# Characterization of the mIF4G domains in the RNA surveillance protein Upf2

**DOI:** 10.1101/2022.12.12.519865

**Authors:** Edgardo M. Colón, Luis A. Haddock, Clarivel Lasalde, Qishan Lin, Juan S. Ramírez-Lugo, Carlos I. González

## Abstract

Thirty percent of all mutations causing human diseases generate mRNAs with premature termination codons (PTCs). Recognition and degradation of these PTC-containing mRNAs is carried out by the mechanism known as nonsense-mediated mRNA decay (NMD). Upf2 is a scaffold protein known to be a central component of the NMD surveillance pathway. It harbors three middle domain of eukaryotic initiation factor 4G (mIF4G) domains in its N-terminal potentially important in regulating the surveillance pathway. In this study, we defined regions within the mIF4G-1 and mIF4G-2 that are required for proper function of NMD and translation termination in *Saccharomyces cerevisiae* Upf2. In addition, we narrowed down the activity of these regions to an aspartic acid (D59) in mIF4G-1 which is important for NMD activity and translation termination accuracy. Taken together, these studies suggest that inherent charged residues within mIF4G-1 of Upf2 play a role in the regulation of the NMD surveillance mechanism in *S. cerevisiae*.

## Introduction

Eukaryotic gene expression is highly regulated at several levels to ensure the proper synthesis of gene products. Among the myriad processes controlling gene expression, there are several mechanisms collectively known as mRNA surveillance that ensure the quality of mRNA molecules. The most studied one is known as nonsense-mediated mRNA surveillance decay (NMD), which recognizes and targets mRNAs containing premature termination codons (PTCs) for degradation, that would otherwise be harmful or toxic to the cell (Gonzalez et al. 2001; Amrani et al. 2006; Wang et al. 2006; Behm-Ansmant et al. 2007; Kervestin and Jacobson 2012; Schweingruber et al. 2013). Besides the regulation of PTC-containing mRNAs, NMD has also been implicated in the regulation of some normal mRNA transcripts (Alonso 2005; Guan et al. 2006; Wittmann et al. 2006; Rebbapragada and Lykke-Andersen 2009; Palacios 2013; Hug et al. 2016; Celik et al. 2017).

Upf1, Upf2, and Upf3 form the core complex of proteins necessary for NMD in eukaryotes. Yeast Upf2 is a 127 kDa acidic protein that accumulates in the perinuclear region of the cytoplasm (Lykke-Andersen et al. 2000; Mendell et al. 2000; Chang et al. 2007). It acts as an adapter protein when Upf1 binds to its C-terminal and Upf3 to its N-terminal (He and Jacobson 1995; He et al. 1997; Kadlec et al. 2004; Chamieh et al. 2008). Various models have been suggested to explain how these factors elicit NMD, which in turn results in the degradation of PTC-containing RNA (Carter et al. 1996; Ruiz-Echevarria et al. 1998; Amrani et al. 2004; Gehring et al. 2005; Lejeune and Maquat 2005; Kervestin et al. 2012). One possibility that has been studied is that Upf1 ATPase and helicase activities are regulated by Upf1 interaction with Upf2. Before interacting with Upf2, Upf1 remains in a closed form that enhances its binding to RNA, which inversely minimizes its ATPase and helicase function (Chamieh et al. 2008; Chakrabarti et al. 2011). When Upf2 binds, through its C-terminal, to the CH domain of Upf1, it causes a conformational change in Upf1 to an open form, resulting in a reduction of its RNA binding activity while enhancing its ATPase and helicase function (Clerici et al. 2009; Chakrabarti et al. 2011). Much emphasis has been given to this N-terminal Upf1-Upf2 C-terminal interaction, while the potential regulatory role of the Upf2 N-terminal has been neglected.

The Upf2 N-terminal harbors three mIF4G domains similar to those found in the eukaryotic initiation factor eIF4G (Aravind and Koonin 2000; Ponting 2000; Clerici et al. 2014; Fourati et al. 2014). In yeast, residues 30-50 found within mIF4G-1 of the Upf2 N-terminal has been shown to be critical for NMD (Wang et al. 2006). Similarly, when human UPF2 mIF4G-1 or mIFG-2 are deleted, individually or together, NMD activity is impaired (Clerici et al. 2014). The crystallized structure of yeast mIF4G-1 revealed that it is formed by 11 □-helices, where 5 pairs are ordered as antiparallel helices stacked against each other (h1-h10) and an additional □-helix (hA) preceding the N-terminal (Fourati et al. 2014). Comparison of human UPF2 mIF4G-1 with yeast mIF4G-1 shows that they are similar, differing only in that 1) the human mIF4G-1 helix hA is much longer, 2) harbors an additional helix (h8i) inserted between helices h8 and h9, 3) has an extra □-helix (hB), and 4) the mIF4G-1 domain structure is more compact (Clerici et al. 2014; Fourati et al. 2014). In yeast this domain harbors phosphorylated residues Ser32 and Ser33 which have been shown to be required for NMD (Wang et al. 2006). Unphosphorylated residues within the yeast mIF4G-1 domain have also been shown to be essential for NMD (Fourati et al. 2014).

In this study, we hypothesized that since the Upf2 mIF4G domains are involved in regulating NMD, then some residues within these domains may be crucial for this activity. To test this hypothesis, Northern Blot analysis and nonsense suppression assays were performed. Results showed a functional region of ten residues withing each one of Upf2 mIF4G-1 and mIF4G-2 domains that are essential for both NMD and translation termination accuracy. Since phosphorylation has been shown to regulate NMD activity, we identified two novel phosphorylated residues in mIF4G-1 domain functional region and one in mIF4G-2 domain functional region. Mutational analysis of the phosphorylated residues did not impair NMD activity.

Further characterization of these domains showed that segments of three and four residues are required for both NMD and translation termination accuracy. Specifically, a mutation of an aspartic acid within one of the segments in the mIF4G-1 impaired the activity of both mechanisms. The results presented here provide an insight for future studies that seek to uncover the exact mechanism by which Upf2 regulates NMD.

## Results

### Identification novel phosphorylation sites in yeast Upf2 mIF4G domains

Previous studies demonstrated that Upf2 is phosphorylated in *S. cerevisiae* and several phosphorylated residues were identified at the mIF4G-1 domain (Wang et al. 2006). To identify novel phosphorylation sites in Upf2 mIF4G domains, protein was purified from a *upf2Δ S. cerevisiae* strain expressing a Flag-tagged UPF2 allele within a plasmid. Purification was performed in the presence of phosphatase inhibitors to avoid the loss of phosphorylation from residues. Purified sample was resolved by 10% SDS-PAGE and stained with Coomassie blue (**Fig 1A**). Flag-Upf2 migrated to approximately the 127 kDa region (**Fig 1A, lane 9**), consistent with previous reports (Cui et al. 1995; Wang et al. 2006). Western blot analysis confirmed that the prominent band observed in the SDS-PAGE was indeed Flag-Upf2 (**Fig 1B**).

**Fig 1.**
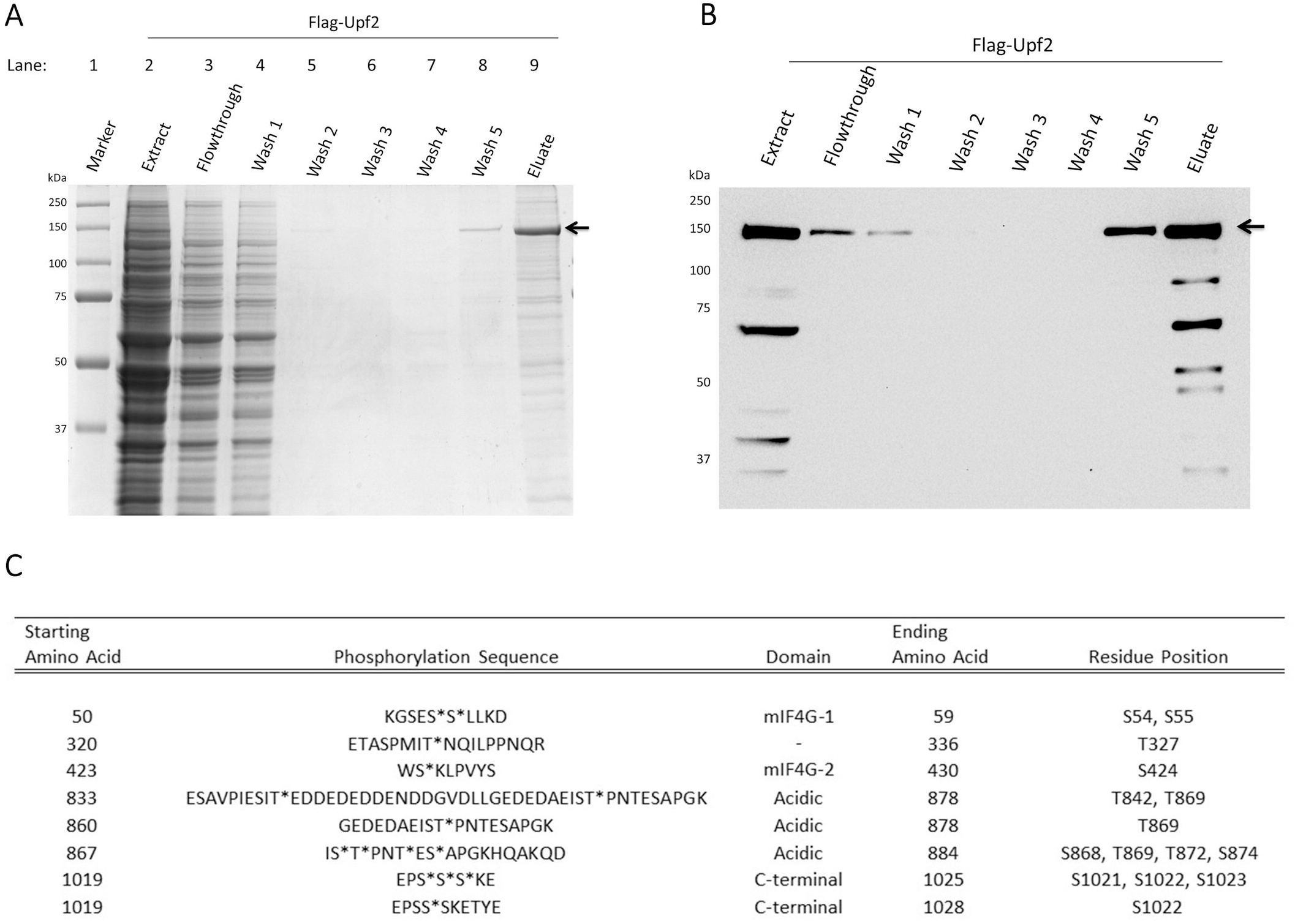
Identification of novel phosphorylation sites in Upf2 mIF4G-1 and mIF4G-2 domains. (A) Coomassie blue-stained 10% SDS-PAGE of immunopurified Flag-Upf2 which is depicted by an arrow (←). (B) Western blot of immunopurified Flag-Upf2 protein depicted by an arrow (←). (C) Trypsin and Gluc-c in-gel digested Upf2 resulting peptides. The resulting peptides were resolved by an ABSCIEX QSTAR XL mass spectrometer (AB Sciex, Framingham, USA) and analyzed as described in Materials and methods. First and fourth columns states the starting and ending residues of the peptide, respectively. Phosphorylated residues in the second column are depicted by an asterisk (*). Domain containing the peptides are specified in the third column. The last column shows the phosphorylated residue position within Upf2. njdsbjds

The Coomassie-stained band harboring Upf2 (~127 kDa) was excised and subjected separately to trypsin and gluc-c digestion. Proteolytic peptides were analyzed by mass spectrometry (see Materials and Methods), which identified eight phosphorylated peptides (**Fig 1C**).

Previously suggested phosphorylated residues S32 and S33 (Wang et al. 2006) were not identified as phosphorylated although various peptide fragments harboring these residues were analyzed. This may have been due to loss of phosphorylation during protein extraction, protein purification, or mass spectrometry analysis of Upf2. Overall, twelve novel residues were identified as phosphorylated in *S. cerevisiae* Upf2, of which three are in Upf2 mIF4G domains: two in the mIF4G-1, and one in the mIF4G-2 (**Figs 1C** and **2A**). A total of 698 of the 1089 amino acids in *S. cerevisiae* Upf2 were present among the analyzed peptides (64% sequence coverage) (**Fig 2B**).

**Fig 2.**
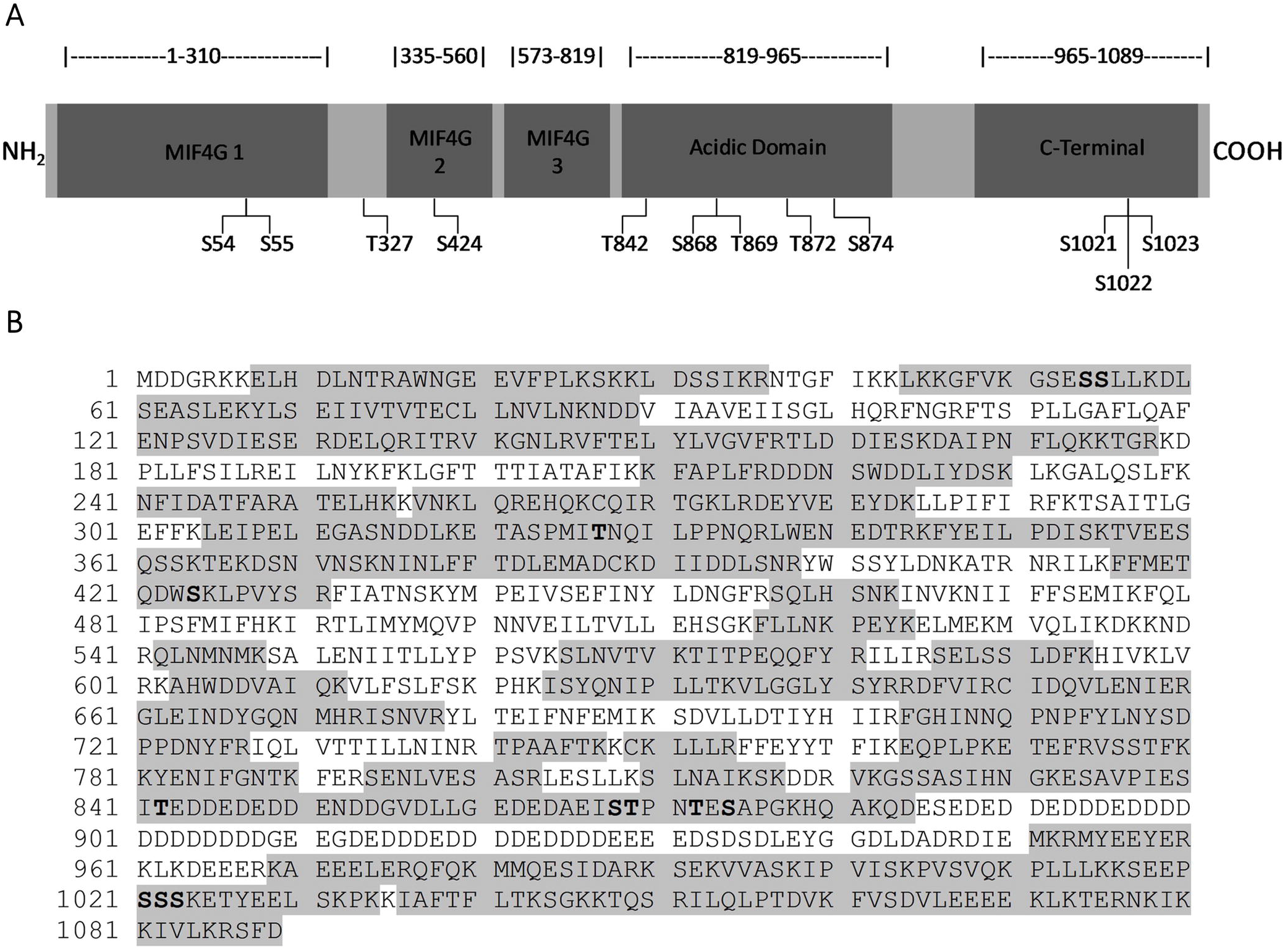
Three phosphorylation sites were identified in Upf2 mIF4G domains. (A) Illustrative representation of Upf2 showing the locations of the MIF4G-1 domain, MIF4G-2 domain, MIF4G-3 domain, acidic domain, and C-Terminal domain. The twelve phosphorylated residues identified by mass spectrometry are presented. (B) *S. cerevisiae* Upf2 amino acid sequence. Sixty-four percent (64%) of sequence coverage was obtained with the mass spectrometry analysis. Analyzed sequences are shown in gray, and phosphorylated residues are in bold.

Sequence alignment showed that none of the Upf2 phosphorylated residues are conserved between species (**Fig 3**). However, phosphorylation of residues S54, S424 and T842 in *S. cerevisiae* showed a tendency to maintain a negative charge at those positions when compared to human UPF2 (**Fig 3**). Residue S54 is located within the mIF4G-1, S424 is located within the mIF4G-2 domain (**Fig 2A**), while residue T842 is in the acidic domain (**Fig 2A**). More importantly, the amino acid D59 (adjacent to the phosphorylated residue in the N-Terminal mIF4G-1 domain) and E843 (adjacent to the phosphorylated residue in the acidic domain) showed a conservation among species of strongly similar properties, namely their acidity given by the negative charge, suggesting an important role for these residues in the biological activity **(Fig 3).** Interestingly, both the mIF4G-1 and acidic domains have been shown to be necessary for NMD activity (**Fig 2A)** (Wang et al. 2006; Fourati et al. 2014).

**Fig 3.**
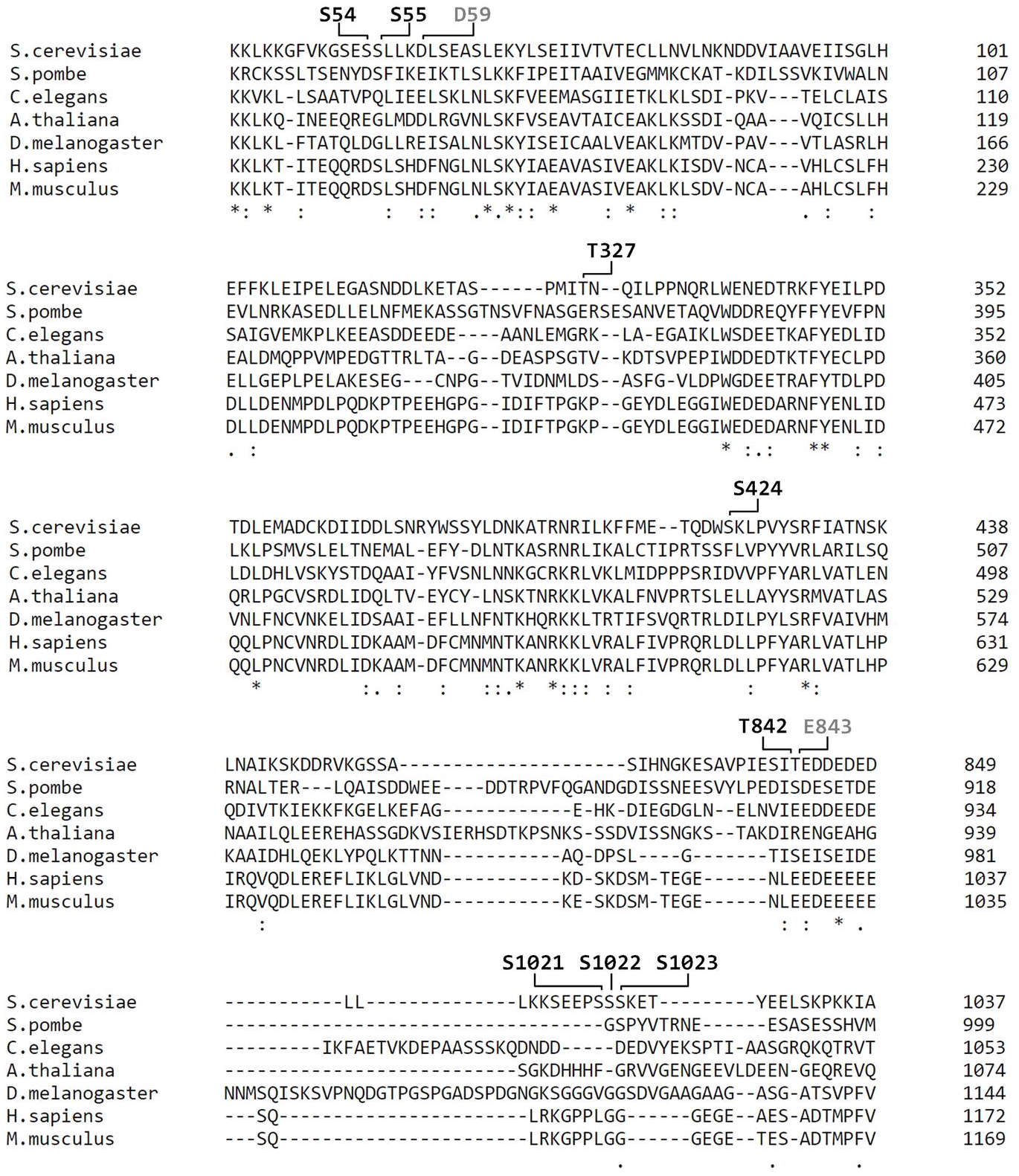
Several residues within Upf2 show a strong conservation of their negative charge. Alignment of sequences from Upf2 *S. cerevisiae* (NCBI NP_011944.2), *S. pombe* (NCBI NP_593784.1), *A. thaliana* (NCBI NP_181459.4), *C. elegans* (NCBI NP_500974.2), *D. melanogaster* (NCBI NP_572434.1), *H. sapiens* (NCBI NP_056357.1), and *M. musculus* (NCBI NP_001074601.1) shows a conservation of the negative charge of residues D59, and E843 as depicted by the colon (:). Phosphorylated residues are depicted in black while nearby negative residues are in gray. Alignment was constructed with Clustal O (version 1.2.4).

### Upf2 mIF4G domain regions harboring phosphorylated residues are required for NMD and translation termination accuracy

To assess the role of Upf2 phosphorylation, the phosphorylated residues were grouped and centered within five different regions. Except for one, each region consisted of ten residues, and were deleted separately (**Fig 4A**). The ‘phospho-regions’ were: 1 (K50-D59), 2 (E321-I330), 3 (Q421-S430), 4 (G831-Q880), and 5 (E1019-E1028). Only phospho-region 1 is located in the mIF4G-1; phosphor-region 2 is located in the region between mIF4G-1 and mIF4G-2; and, phospho-region 3 is located within the mIF4G-2 domain (Wang et al. 2006; Kervestin and Jacobson 2012) (**Fig 4A**).

**Fig 4.**
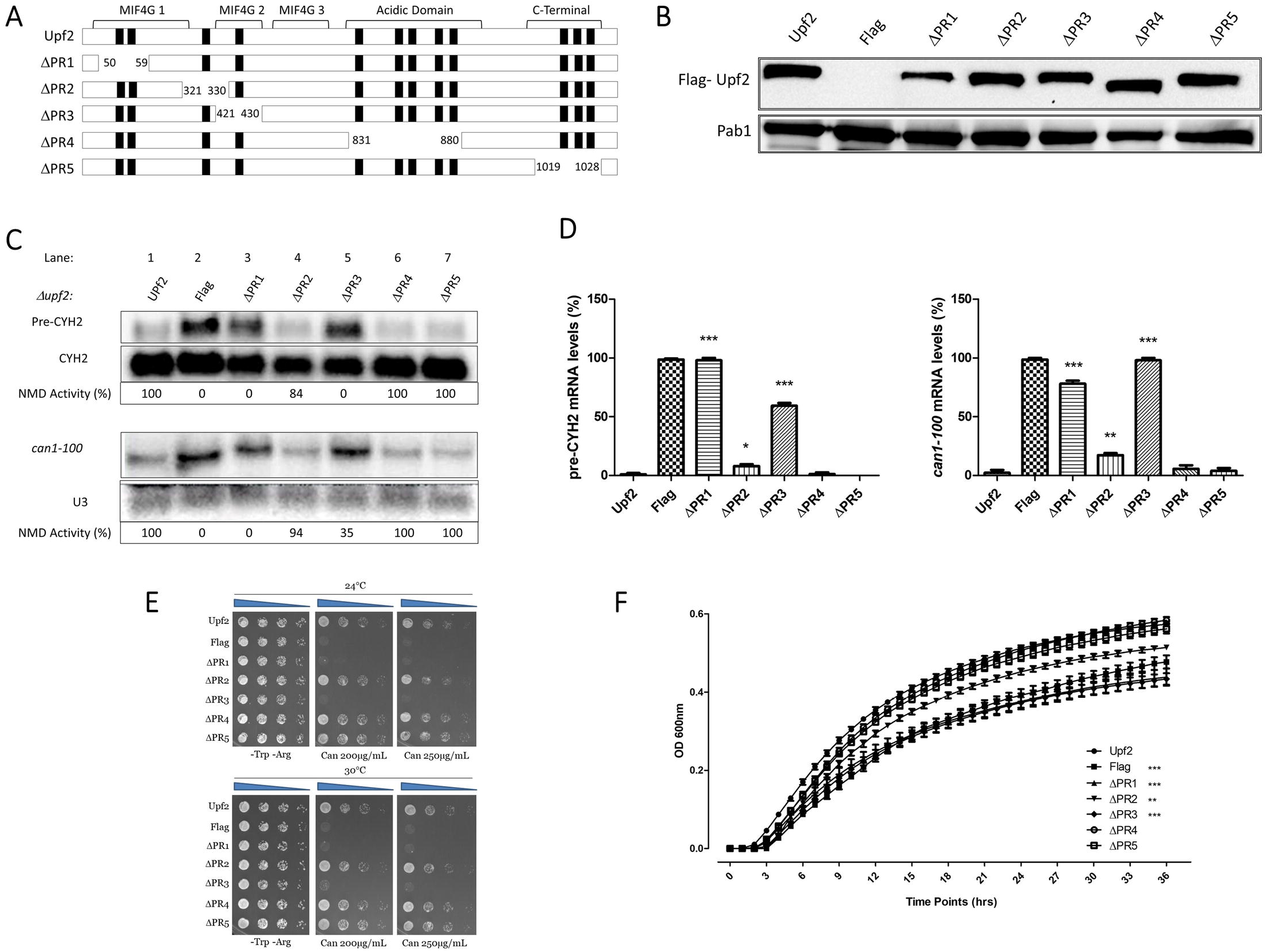
NMD activity requires Upf2 mIF4G-1 and mIF4G-2 phosphorylated regions 1 and 3. (A) Illustrative representation of Upf2 phospho-region mutants (ΔPR). Phosphorylated residues are depicted as black rectangles. In the phospho-region 1 (ΔPR1) residues K50-D59 were deleted, in phospho-region 2 (ΔPR2) residues E321-I330 were deleted, phospho-region 3 (ΔPR3) residues Q421-S430, in phospho-region 4 (ΔPR4) residues G831-Q880 were removed, and in phospho-region 5 (ΔPR5) residues E1019-E1028 were removed. (B) Analysis of cytoplasmic extracts by Western blot showing Upf2 expression. Poly(A) binding protein (Pab1) was used as a loading control. (C) Northern blot from total cellular RNA was used to determine NMD activity of the Upf2 phospho-region deletions. (D) *Pre-CYH2* mRNA accumulation from Northern blot expressed as a mean value ± standard deviation. (E) *can1-100* nonsense suppression assay was used to assess the role of Upf2 in translation termination efficiency. Wild-type and mutant *upf2* yeast strains were serially diluted (1:10) five times. F) *can1-100* nonsense suppression assay growth curves. Two-way ANOVA was used for statistical analysis. Significant results (P<0.05) when compared to the WT strain are depicted with asterisks (*).

Western blot analysis confirmed the expression of these *upf2* mutants (**Fig 4B**). Northern blot analysis was used to determine NMD activity of these mutant strains by detecting the premature termination codon containing transcripts *CYH2* pre-mRNA and *can1-100* mRNA. *CYH2* pre-mRNA contains the in-frame premature termination codon (He et al. 1993), which in conjunction with the mature *CYH2* mRNA provides a ratio that has been routinely used as a measurement of NMD activity (He et al. 1993; Wang et al. 2006; Lasalde et al. 2014). Levels of *can1-100* were measured by normalizing against the mRNA loading control U3. Results from total cellular RNA analysis from WT and phospho-region 2 (E321-I330), 4 (G831-Q880), and 5 (E1019-E1028) mutant strains showed that NMD activity was not affected in comparison to the defect observed in the *upf2Δ* strain (**Figs 4C and 4D)**, compare lane 1 with lanes 4, 6, 7). This demonstrated that none of these deletions affected NMD activity. But deletion of phospho-region 1 (K50-D59) and phospho-region 3 (Q421-S430) showed reduced NMD activity like that of the *upf2Δ* strain, demonstrating that these deletions affected NMD activity (**Figs 4C and 4D**, lane 3 and 5). Phospho-region 1 of Upf2 harbors the phosphorylated residues S54 and S-55, while phospho-region 3 of Upf2 harbors the phosphorylated residue S424 (**Fig 3**).

In addition to being critical for NMD, Upf2 also promotes translation termination (Maderazo et al. 2000). The *can1-100* nonsense suppression assay was used to qualitatively and quantitatively assess the role of Upf2 phospho-region 1 and phospho-region 3 in translation termination efficiency. This assay exploits the fact that in a *can1-100* strain defective in translation termination, read-through of the nonsense codon produces a functional form of the protein Can1, an arginine permease capable of importing the toxic arginine analog canavanine. Consequently, lower growth rates of the different Upf2 mutant strains in comparison to the WT are used to measure enhanced nonsense suppression activity (Ono et al. 1983; Maderazo et al. 2000; Estrella et al. 2009). Results from the *can1-100* nonsense suppression assay demonstrated that, similar to the wild-type UPF2 strain, strains with deletions of phospho-region 2, phospho-region 4, and phospho-region 5 grow normally and thus do not show a nonsense suppression phenotype (**Fig 4E**). In contrast, strains harboring the deletion of phospho-region 1 and phospho-region 3 show less growth in the presence of canavanine, suggesting a reduction in translation termination efficiency (**Fig 4E**). In addition, strains with deleted phospho-region 1 and phospho-region 3 grown in synthetic complete liquid media with canavanine over a period of time show decreased growth, also indicating that these phospho-regions play a role in translation termination efficiency (**Fig 4F**). Taken together, these data suggest that two small regions of Upf2 consisting of ten residues and harboring phosphorylated residues, are critical for NMD activity as well as a proper translation termination.

### Upf2 mIF4G-1 S54 and S55, and mIF4G-2 S424, phosphorylated residues are not required for NMD and translation termination accuracy

Examination of phospho-region 1 showed a serine, S52 **(Fig. 4A)**, adjacent to the serine we found to be phosphorylated in this study, S54 and S55 (**Fig 2A**). Likewise, examination of phospho-region 3 showed a tyrosine and serine, Y429 and S430 **(Fig 4A)**, adjacent to the phosphorylated residue S424 identified here (**Fig 2A**). These adjacent residues were not identified as a phosphorylated residues in our mass spectrometry analysis (**Fig 2A**). Nonetheless, these amino acid residues may potentially be phosphorylated under different conditions or phosphorylated at levels in which our analysis cannot detect (**Fig 1D**). To test the role of the phosphorylated residues identified and adjacent residues, single, double, and triple mutants were generated using site-directed mutagenesis (**Fig 5A**).

**Fig 5.**
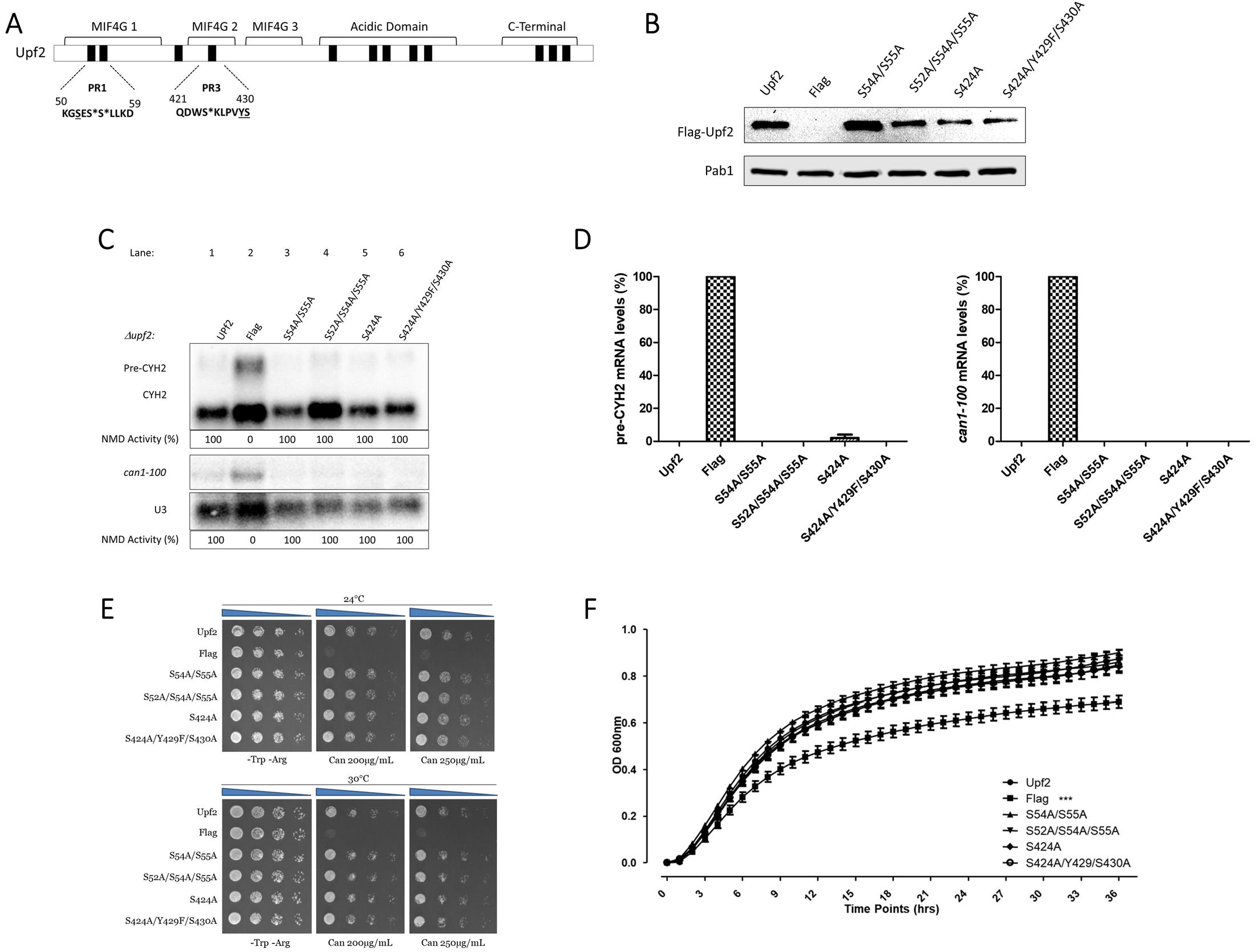
NMD activity does not require Upf2 mIF4G-1 and mIF4G-2 phosphorylated residues of either phospho-region 1 or phospho-region 3. (A) Illustrative representation of Upf2 phospho-region (PR) mutants 1 and 3. Phosphorylated residues S54, S55 and S424 are depicted with an asterisk (*) while adjacent potential phosphorylated residues S52, Y429, S430 are underlined. (B) Analysis of cytoplasmic extracts by Western blot showing Upf2 expression. Poly(A) binding protein (Pab1) was used as a loading control. (C) Northern blot from total cellular RNA was used to determine NMD activity of the Upf2 phosphorylated residues substitutions. (D) *pre-CYH2* mRNA accumulation from Northern blot expressed as a mean value ± standard deviation. (E) *can1-100* nonsense suppression assay was used to assess the role of Upf2 in translation termination efficiency. Qualitatively wild-type and mutant *upf2* yeast strains were serially diluted (1:10) five times. (F) *can1-100* nonsense suppression assay growth curves. Two-way ANOVA was used for statistical analysis. Significant results (P<0.05) when compared to the WT strain are depicted with asterisks (*).

The serine and tyrosine residues were mutated to alanine and phenylalanine, respectively, to mimic unphosphorylated versions of these amino acids. One single point mutant (S424A), one double mutant (S54A/S55A), and two triple mutants (S52A/S54A/S55A and S424/Y429F/S430) were generated. Expression of these *upf2* mutants was confirmed by Western blotting (**Fig 5B**). Northern blot analysis of total cellular RNA isolated from WT and the five mutant strains were used to determine NMD activity as described previously. As anticipated, NMD activity in the wild-type strain was 100%, while no activity was observed in the *upf2Δ* strain (**Fig 5C**, compare lanes 1 and 2). All the mutant constructs retain NMD activity at a level similar to the WT (**Figs 5C and 5D**, compare lane 1 with lanes 3, 4, 5 and 6). Likewise, none of the mutant strains showed a decrease in growth in the presence of canavanine (**Figs 5E and 5F**). Taken together, these data suggest that both NMD activity and translation termination accuracy are independent of the phosphorylation of S52, S54, and S424. In addition, nearby residues S52, Y429 and S430 do not have a redundant phosphorylation role in the absence of phosphorylated residues S54, S55, and S424, and are also not essential for NMD activity and translation termination.

### Segments within Upf2 mIF4G-1 and mIF4G-2 are essential for NMD and translation termination accuracy

Activity of the studied regions appeared to be independent of their phosphorylation status. But, since the absence of these regions diminishes NMD activity, which might be due to impairments in the structure of Upf2, we sought to further characterize them by studying other segments within them. To assess their role in NMD, two segments containing up to a maximum of four residues were deleted from each region. The deleted segments were: 1 (K50-E53), 2 (L57-D59) from phospho-region 1 in mIF4G-1, and 3 (Q421-W423), 4 (K425-V428) from phospho-region 3 in mIF4G-2. Constructs harboring these deletions were transformed into an *upf2Δ* strain (**Fig 6A**). Expression of these *upf2* mutants was confirmed by Western blotting (**Fig 6B**). The WT and mutant strains were subjected to Northern blot analysis to determine the NMD activity as described above. Levels of *pre-CYH2* mRNA showed a reduction of NMD activity of 64%, 63% and 66% for mutants lacking residues L57-D59, Q421-W423 and 425-428, respectively, and a reduction of 84%, 54% and 45%, respectively, when *can1-100* mRNA levels were measured (**Figs 6C and 6D,** lane 4-6). In contrast, deletion of residues K50-E53 had no significant effect on NMD activity (**Figs 6C and 6D,** lane 3). Results from the *can1-100* nonsense suppression assays show that deletion of segment 1 (K50-E53) did not present a nonsense suppression phenotype, similar to the wild-type UPF2 strain (**Figs 6E and 6F**). In contrast, strains harboring the deletion of segment 2 (L57-D59), segment 3 (Q421-W423), and segment 4 (K425-V428) exhibited reduced growth in the presence of canavanine, pointing to a decrease in translation termination efficiency (**Figs 6E and 6F**). All together, these data suggest that the regulation of NMD is modulated by residues residing within segment 2 (L57-D59) of mIF4G-1, and within segments 3 (Q421-W423) and 4 (K425-V428) of mIF4G-2.

**Fig 6.**
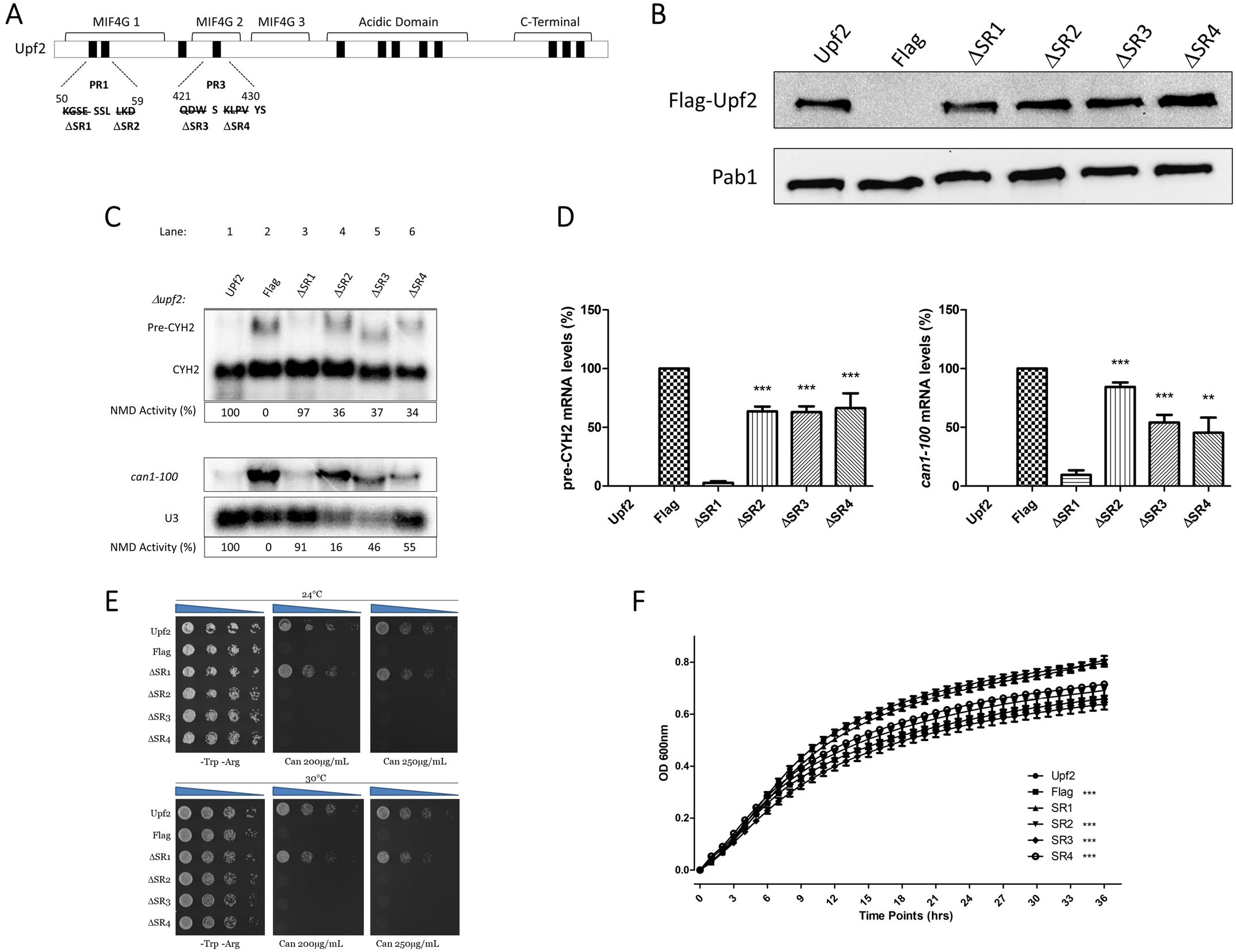
NMD activity requires Upf2 mIF4G-1 and mIF4G-2 non-phosphorylated segments. (A) Illustrative representation of Upf2 non-phosphorylated segments (SR) within phospho-region (PR) 1 and 3. Depicted by a strikethrough, in the segment 1 (SR1) residues K50-E53 were deleted, in segment 2 (SR2) residues L57-D59 were deleted, segment 3 (SR3) residues Q421-W423, and in segment 4 (SR4) residues K425-V428 were removed. (B) Analysis of cytoplasmic extracts by Western blot showing Upf2 expression. Poly(A) binding protein (Pab1) was used as a loading control. (C) Northern blot from total cellular RNA was used to determine NMD activity of the Upf2 non-phosphorylated segments deletions. (D) *pre-CYH2* mRNA accumulation from Northern blot expressed as a mean value ± standard deviation. (E) *can1-100* nonsense suppression assay was used to assess the role of Upf2 in translation termination efficiency. Wild-type and mutant *upf2* yeast strains were serially diluted (1:10) five times. (F) *can1-100* nonsense suppression assay growth curves. Two-way ANOVA was used for statistical analysis. Significant results (P<0.05) when compared to the WT strain are depicted with asterisks (*).

### Upf2 mIF4G-1 residue D59 is required for NMD and translation termination accuracy

Effects seen on NMD regulation when segment 2, segment 3 and segment 4 are deleted might still be due to an impairment of Upf2 structure. To discard this possibility, we tested the role of the residues within these segments of mIF4G-1 and mIF4G-2 through single mutants generated using site-directed mutagenesis (**Fig 7A**). Charged residues were mutated to incorporate charge inversions. Negatively charged residues D59 and D422 were substituted by lysine generating D59K and D422K. The positively charged residue K425 was substituted by glutamic acid generating K425E, while neutral residue W423 was substituted by alanine generating W423A. Expression of these mutants was confirmed by Western blotting (**Fig 7B**). Total cellular RNA isolated from WT and the five different point mutant strains was analyzed by Northern blot and NMD activity was ascertained as described above. Levels of *pre-CYH2* mRNA showed a 20% reduction of NMD activity for the point mutant D59K while a 15% reduction when *can1-100* mRNA levels were measured (**Figs 7C and 7D**, lane 3). In contrast, point mutants D422K, W423A and K425E had no significant effect on NMD activity. (**Figs 7C and 7D**, lane 4-6). Likewise, the point mutant D59K showed a decrease in growth when grown in the presence of canavanine (**Fig 7E**). Taken together, these data suggest that the regulation of NMD is influenced by the residue D59 residing within the mIF4G-1 of Upf2.

**Fig 7.**
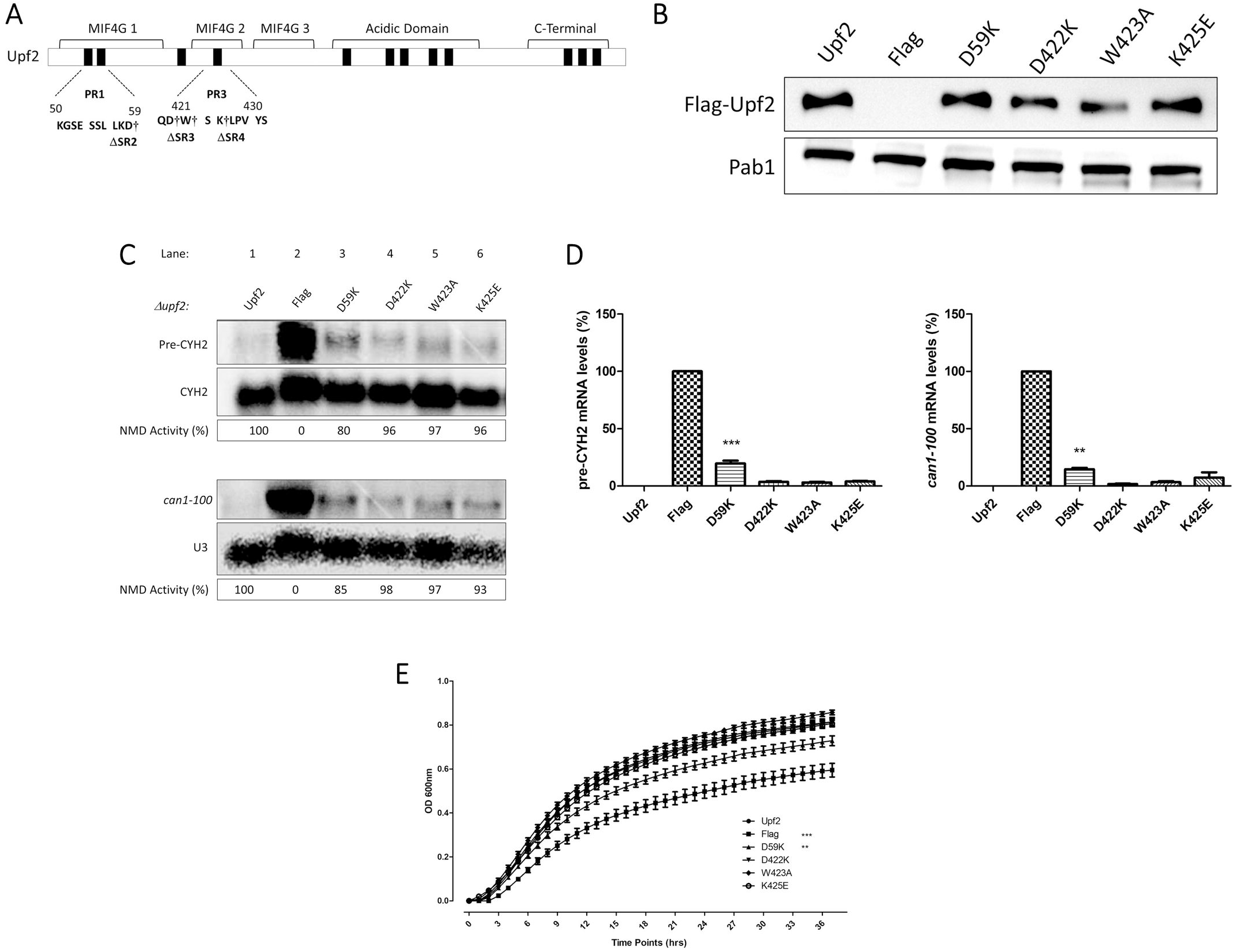
NMD activity requires Upf2 mIF4G-1 residue D59. (A) Illustrative representation of Upf2 sub-region mutants. Residues D59, D422, W423, and K425, which were substituted, are depicted with a dagger (†). (B) Analysis of cytoplasmic extracts by Western blot showing Upf2 expression. Poly(A) binding protein (Pab1) was used as a loading control. (C) Northern blot from total cellular RNA was used to determine NMD activity of the Upf2 single mutant deletions. (D) *pre-CYH2* mRNA accumulation from Northern blot expressed as a mean value ± standard deviation. (E) *can1-100* nonsense suppression assay growth curves. Two-way ANOVA was used for statistical analysis. Significant results (P<0.05) when compared to the WT strain are depicted with asterisks (*).

## Discussion

Most studies of the regulation of NMD has been focused on Upf1 enzymatic properties (Page et al. 1999; Yamashita et al. 2001; Grimson et al. 2004; Guan et al. 2006; Wang et al. 2006; Okada-Katsuhata et al. 2012; Kerenyi et al. 2013; Lasalde et al. 2014), in comparison to Upf2 (Chiu et al. 2003; Wang et al. 2006; Clerici et al. 2014). One distinct research area that has been neglected is Upf2 protein-protein interactions, mediated by its mIF4G domains, which serve as a mean for regulating NMD. In *S. cerevisiae*, Upf3 has been shown to interact with Upf2 mIF4G-3 and abolishment of this interaction eliminates NMD activity (He et al. 1997; Serin et al. 2001). The mIF4G-1 domain is also involved in mediating protein-protein interactions where Hrp1 has been shown to bind to this domain (Gonzalez et al. 2000; Wang et al. 2006) and disruption of this interaction decreases the activity of NMD (Wang et al. 2006).

In contrast, human UPF2 mIF4G-3 interacts with the Upf3 RNA recognition motif (RRM) via charged residues that, when mutated, abolish the Upf2-Upf3b interaction, leading to inhibition of NMD (He et al. 1997; Kadlec et al. 2004; Kashima et al. 2006). Similar to Upf3 binding, Upf2 mIF4G-3 can also interact with RNA, and mutations R796E and R797E impair Upf2 mIF4G-3 RNA binding activity (Kadlec et al. 2004). SMG1 has also been shown to interact with Upf2 through its mIF4G-3 domain (Kashima et al. 2006; Clerici et al. 2014). This interaction, as well as the Upf2-Upf3 interaction, are necessary for the association of SMG-1 with the Y14-post-splicing RNA complex to elicit phosphorylation of Upf1 (Kashima et al. 2006). Interaction of SMG1 and Upf2 allows the phosphorylation of the S1046 residue located in the mIF4G-3, although this phosphorylation does not inhibit NMD activity (Clerici et al. 2014). eRF3 has been previously shown to interact with Upf2 mIF4G-3 less strongly than Upf3 (Lopez-Perrote et al. 2016).

In this article we have shown that there are two regions consisting of ten residues each (residues 50-59 and 421-430) harbored within Upf2 mIF4G-1 and mIF4G-2, respectively, which are essential for proper function of NMD (**Figs 4C and 4D**). Since deletion of ten residues might influence Upf2 structure, and by so affect it function, we reduced the possibility of this being the case by generating smaller deletions which are less likely to have a structural effect. Here we demonstrated that deletion of segments with a minimum of three residues within both mIF4G-1 and mIF4G-2 are sufficient to reduce the activity of NMD (**Figs 6C and 6D**). Most importantly, trough charge inversion mutation analysis, we identified residue D59, present in mIF4G-1, as necessary for activity of NMD (**Figs 7C and 7D**). This residue is located in helix 2 of the mIF4G-1, a region enriched in basic residues as shown in previous studies (Fourati et al. 2014). Our results are consistent with these studies, in which charge inversion of residues K35E/R36E and K42E/K43E/K45E, in helix 1, and K67A/E71A, in helix 3, caused an inhibition of NMD activity. Although mutation of D59 significantly reduced NMD activity, it did not reduce the activity to the same extent as deleting three residues. This may be because deletion of these three residues completely removed the charges present, while the charges were only partially reduced in our D59 mutant. Since deletion of segment 2 which contains the residues L57, K58 and D59 might be affecting protein structure and as a result NMD activity, future experiments should focus on a triple substitution mutant of these residues to discard that possibility. In addition, it remains of interest if charge inversion of residue K58 by itself would have a similar effect on NMD as our D59 mutant and the effect that might have a double charge inversion mutant of K58 and D59. Moreover, D59 showed strong conservation of its negative charge when compared with other eukaryotes **(Fig 3)**. We suspect that the evolutionary conservation of charged residue present at the mIF4G domains might be dictating the specificity of interactions between Upf2 and yet unidentified proteins involved in NMD. Further analysis of *S. cerevisiae* Upf2 mIF4G domains protein-protein interactions are needed to understand their regulatory role in NMD.

Furthermore, we have identified twelve phosphorylated residues in *S. cerevisiae* Upf2 out of which two are in the mIF4G-1 (S54 and S55) and one in the mIF4G-2 (S424) (**Figs 2A and 2B**). Our mutation analysis showed that phosphorylated residues within these domains do not disrupt NMD activity (**Figs 5C and 5D**). These results are consistent with previous findings where, in mammalian cells, Upf2 is phosphorylated in the mIF4G-3 at residue S1046 by SMG-1 but it is not necessary for NMD activity (Clerici et al. 2014). We speculate that Upf2 might be regulated by a complex phosphorylation code where combinations of specific phosphorylated residues enable protein function. Furthermore, our experiments are solely focused on Upf2 phosphorylation as a regulator of NMD mediating degradation of PTC-containing mRNAs. Rather than mediating degradation of PTC-containing mRNAs, Upf2 phosphorylation might play a role in the regulation of normal mRNA transcripts, as has been previously shown (Alonso 2005; Guan et al. 2006; Wittmann et al. 2006; Rebbapragada and Lykke-Andersen 2009; Palacios 2013; Hug et al. 2016). It is possible that specific phosphorylated residues are necessary to elicit the targeting and degradation of PTC-containing mRNAs while others are required for targeting normal mRNAs. Further biochemical analysis should focus on determining cellular events that might trigger the modification of specific residues and their overall function. Because of the regulatory role that phosphorylation exerts over proteins, it remains of interest to determine if there are other phosphorylated residues in *S. cerevisiae* Upf2 which might be involved in NMD.

Besides NMD, in mammals and yeast, Upf2 promotes proper translation termination by repressing read-through of the premature termination codons (Maderazo et al. 2000; Wang et al. 2001). Our results reveal a role for Upf2 mIF4G-1 and mIF4G-2 in translation termination accuracy, suggesting that this event also depends on these regions (**Figs 4 and 6**). In addition, we showed that residue D59 located in the mIF4G-1, which plays a role in NMD, is also involved in the fidelity of translation termination (**Fig 7E**). Hence, we propose that disrupting the evolutionary conserved charged residues in these domains interferes with their ability to interaction with protein necessary for translation termination.

Herein we characterized Upf2 mIF4G-1 and mIF4G-2 domains providing evidence regarding their role in NMD and translation termination accuracy **(Fig 8)**. Characterization of mIF4G-1 proved to be of particular importance since we were able to identify a region of ten residues, a segment of three residues (L57-D59), and a single residue (D59) capable of causing a significant reduction in the activity of NMD. Further biochemical and structural analysis focusing on charges in the mIF4G domains of Upf2 are required to better understand the role of these key regulatory components of the nonsense-mediated mRNA decay pathway.

**Fig 8.**
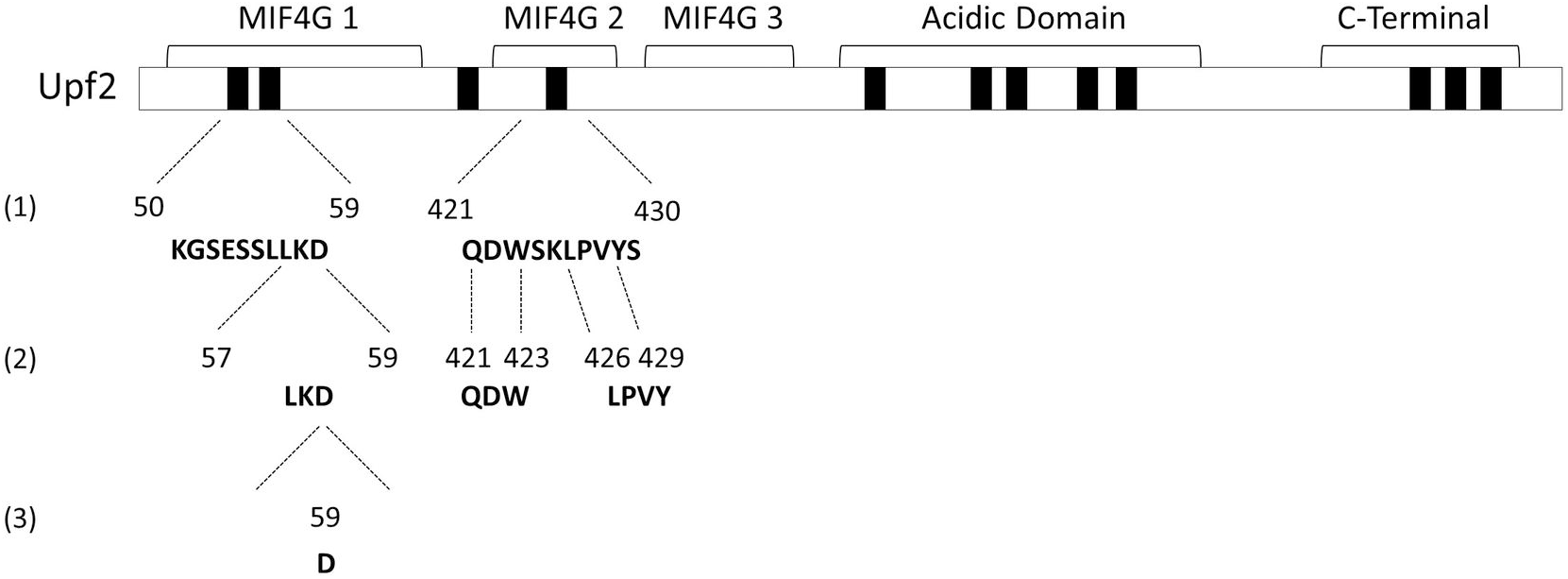
Characterization of *S. cerevisiae* Upf2 mIF4G domains. NMD activity requires Upf2 regions, segments and single residue within mIF4G-1 and mIF4G-2. Schematic representation of Upf2 mIF4G domains characterization. mIF4G-1 and mIF4G-2 regions and segments required for NMD are aligned with the numbers (1) and (2), respectively. Single residue within mIF4G-1 required for NMD is aligned with the number (3).

## Materials and methods

### Strains and plasmids

*S. cerevisiae* strain S288C derivative *upf2Δ (Mat □, ura3-52, trp1-Δ1, leu2-2, tyr7-1, can1-100, his3-Δ200*, UPF1, *upf2Δ1:* HIS3, UPF3) was used as the wild-type strain for all the experiments performed in these studies. The yeast *2μ* plasmid pG-1, containing the Flag-UPF2 allele was used as the vector (Wang and Wilkinson 2001). Briefly, the GAL1 promoter was inserted in pUC8. The promoter contains the regulatory sequence and the transcription initiation site, but not the initiation codon. N-terminal FLAG *2μ* TRP1 pG-1. AMP^R^ was cloned in frame and upstream of the UPF2 allele. Deletions of *upf2* were constructed using the Flag-UPF2 allele as a template, subjected to the single-step cloning method (Makarova et al. 2000). Point mutants of *upf2* were constructed using Flag-UPF2 allele as a template, subjected to the site-directed mutagenesis method (Wang and Wilkinson 2001). Constructs were sequenced and mutations were confirmed by sequence alignment. Yeast strains were transformed by an improved version of the lithium acetate method (Gietz et al. 1992).

### Whole cell protein extract and Western blot

Cells were cultured to an optical density (OD_600_) of 1.0-1.2 followed by centrifugation at 3,200 rpm for 10 minutes at 4°C. Cells were then washed in: 1) 15mL of cold ddH2O and centrifuged; 2) 15mL of cold Buffer XA (20mM HEPES, NaOH pH=7.4, 150mM NaCl, 2.0mM EDTA, 0.01% Triton X-100, 30mM Sodium Fluoride, 30mM β-glycerol phosphate, 5mM Sodium Pyrophosphate, 1mM PMSF, 1X Protease Inhibitor Cocktail, 100nM Okadaic Acid) and centrifuged; and 3) re-suspended in 5mL of Buffer XA. Cells were then centrifuged, followed by the addition of 350-400uL buffer XA and an equal volume of acid-washed glass beads. Vortex was then applied 13 times for 30 seconds with 45 seconds of cooling between steps. Centrifugation was applied for 30 minutes at 13,200 rpm and the lysate containing the protein extract was removed to a new tube. Bovine serum albumin (BSA) was used as a protein standard to quantify total protein extract. Proteins were resolved in 10% SDS-PAGE and transferred to a nitrocellulose membrane (Bio-Rad). To detect Flag-Upf2, anti-Flag (Sigma) and anti-Mouse peroxidase conjugated (Sigma) were used as primary and as secondary antibody, respectively. Pab1 was used as a loading control. Exposure of the membranes was performed using SuperSignal West Dura chemiluminescent substrate (Thermo Scientific) in a Molecular Imager^®^ ChemiDoc™ XRS+ with Image Lab™ Software (BioRad).

### Upf2 protein purification

Upf2 was purified using anti-Flag M2 affinity gel (Sigma), as previously described (Wang et al. 2006). Briefly, cell protein extract was performed as described above. Propylene chromatography columns were filled with 1mL of anti-Flag M2 affinity gel (Sigma). 15mL of 1X TBS-T were used twice to wash the beads and 1mL-original volume of Buffer XB (20mM HEPES, NaOH pH=7.4, 150mM NaCl, 2.0mM EDTA, 0.01% Triton X-100, 30mM Sodium Fluoride, 30mM β-glycerol phosphate, 5mM Sodium Pyrophosphate, 1mM PMSF) was added to the anti-Flag beads. The beads were then added to the protein extracts in the column and incubated overnight at 4°C with constant rotation. Afterwards, the beads were washed once with 3mL of Buffer XB, three times with Buffer XB-2 (20mM HEPES, NaOH pH=7.4, 300mM NaCl, 2.0mM EDTA, 0.01% Triton X-100, 30mM Sodium Fluoride, 30mM β-glycerol phosphate, 5mM Sodium Pyrophosphate, 1mM PMSF) and lastly four times with the Buffer XB. Beads were then incubated with 200 ul of Buffer XC (20mM HEPES, NaOH pH=7.4, 150mM NaCl, 2.0mM EDTA, 0.01% Triton X-100, 30mM Sodium Fluoride, 30mM β-glycerol phosphate, 5mM Sodium Pyrophosphate, 1mM PMSF, Flag Peptide 5ug/ul) for 15 minutes at 4°C. Elution 1 was collected followed by the addition of 300ul of Buffer XC to the beads to collect elution 2. Finally, 300ul of Buffer XC was added to the beads to collect elution 3. Elutions were resolved in 10% SDS-PAGE, stained with Coomassie Brilliant Blue (Sigma) and analyzed by Western blot. BSA was used as a protein standard to determine the protein concentration.

### Tandem mass spectrometry analysis

Prior to analysis, sample gel pieces from three different transformants expressing the Flag-UPF2 allele were washed, reduced, alkylated, and in-gel tryptic digested. Additional analysis was be performed by in-gel gluc-c digestion. Proteolytic peptides were extracted from the gel. The phosphopeptides were enriched by TiO_2_ TopTip (Glygen, Inc). Both flow-through and elute was analyzed by LC-MS/MS, respectively. For this analysis, an ABSCIEX QSTAR XL mass spectrometer (AB Sciex, Framingham, USA) was used. This MS was coupled to a CapLC (Waters Co. Milford, MA, USA) HPLC with a Phenomenex C18 (3μm, 300A, 100μm ID × 150mm, Phenomenex, Torrence, CA) analytical column, and a Vydac Everest C18, 300A, 5μm trap column. The solvents used were composed of 5 % CH_3_CN + 0.1% formic acid + 0.01% TFA (Solvent A) and 85% CH_3_CN + 10% isopropanol + 5% H_2_O + 0.1% formic acid + 0.01% TFA (Solvent B). Samples were kept at a 250nl/min flow rate and a 60 min linear gradient from 10% to 100% B.

### Mass spectrometry data analysis

MS/MS spectra files were processed and combined using Mascot distiller software from the MatrixScience with a processing macro that smooths, centers, and assesses the quality of data. In-house MASCOT 2.3 from Matrix Science (London, UK) was used to assist the interpretation of tandem mass spectra against targeted protein sequences. Error Tolerant Search was used in the analysis. Spectra that remained unmatched which contained adequate information were subjected to an Error Tolerant Search. All database entries from a first pass search, that contained one or more peptide matches with scores at or above the homology threshold, were selected for an error tolerant, second pass search. In this second pass search the complete list of modifications was tested. The list of modifications used by Mascot was taken directly from the Unimod database. The entries on the modification list were tested serially, and all permutations of each individual modification were analyzed. Mass delta of the modification were rejected if they were less than the smaller of the precursor mass tolerance and the fragment mass tolerance.

### RNA isolation and Northern blot

Total RNA was isolated using the hot phenol method (Herrick et al. 1990). Random-primed DNA probes were prepared from a 0.6-kb EcoRI-HindIII fragment spanning a region of the *CYH2* mRNA. Northern blots were quantified using a BioRad Molecular Imager FX. The activity of NMD was calculated by comparing the ratio of *pre-CYH2* to *CYH2* in *Δupf2* strains transformed with either a vector (0%) or wild-type Upf2 (100%). Values of mRNA level represent averages ± standard deviation from three independent experiments.

### *can1-100* nonsense suppression assay

Canavanine drug sensitivity test was performed as previously described (Maderazo et al. 2000). Briefly, wild-type and mutant strains were grown in synthetic liquid media lacking arginine and tryptophan to mid-log phase (OD_600_ = 0.8-0.9). Samples from these cultures were serially diluted (1:10) five times, and aliquots of the four dilutions were spotted on synthetic plates lacking arginine and without or with 200μg and 250μg of canavanine. Plates were incubated at 24°C and 30°C for 48 hr.

### *can1-100* nonsense suppression assay growth curves

Growth curves were performed using a Synergy H1 Hybrid Multi-Mode Microplate Reader (BioTek Instruments, Winooski, VT). Briefly, wild-type and mutant strains were grown in synthetic liquid medium lacking arginine and tryptophan to mid-log phase at 30°C and diluted to an OD_600_ = 1.6. From there, cells were plated into a Falcon 96 well clear flat bottom untreated cell culture microplate (Corning) at a final OD_600_ = 0.25, and 500μg of canavanine were added. Measurements were taken every hour, with an initial 0 hr reading, for thirty-six hours under continuous high orbital shaking. Shaking speed was set to fast and frequency set to 559 (1mm amplitude). Temperature was set at 30°C. Cell growth was assessed by absorbance measurements made at 600nm. Reads were collected using Gen5™ Data Analysis Software (BioTek Instruments, Winooski, VT). Growth curves were generated using GraphPad.

## Acknowledgements

We are grateful to Dr. Michael Culbertson (University of Wisconsin-Madison Emeritus Professor) for sharing with us the *S. cerevisiae* strain S288C derivative *upf2Δ*. We are thankful to the DNA Sequencing and Genotyping Facility at the University of Puerto Rico-Rio Piedras Campus (UPR-RP) for their assistance. We are very appreciative of the effort from all the members of the González laboratory.

